# An application of fNIRS for the prediction of hearing thresholds in sleeping infants: a proof of principle

**DOI:** 10.64898/2026.11.09.764747

**Authors:** D. X. Gao, D. Mao, J. Wunderlich, J. Esmaelpoor, G. Balasubramanian, C. M. McKay

## Abstract

This study applied a two-component-response modelling approach to fNIRS recordings obtained from naturally sleeping infants presented with auditory stimuli at different sound intensities. We investigated whether the relationship between fNIRS response measures and stimulus intensity could be modelled to estimate individual auditory hearing thresholds. In the normal-hearing group, 75% of infants had threshold estimates for the sound /ba/ within the expected normal-hearing range, with an estimated mean equivalent normal loudness (ENL) of 10.72 dB HL (SD 8.64 dB HL). A case study further explored the potential application of this approach in an infant with hearing impairment, highlighting the importance of reliable responses at lower stimulus intensities and appropriate regression model selection. These findings provide preliminary evidence that cortical haemodynamic responses can be quantitatively related to sound intensity for estimating auditory sensitivity.

## 1 Introduction

Accurate assessment of hearing in early infancy is essential for identifying hearing impairment and supporting timely intervention. Objective physiological measures, including auditory brainstem response (ABR) (Jewett and Williston, 1971) and otoacoustic emissions (OAEs) (Kemp, 1978), are widely used for hearing assessment when reliable behavioural responses cannot yet be obtained. However, these measures primarily assess peripheral and subcortical auditory function and do not directly measure cortical responses to sound. Additionally, ABR cannot be used to estimate hearing thresholds in infants diagnosed with auditory neuropathy, leading to detrimental delays in intervention for these infants. An objective method that is capable of measuring auditory cortical responses could provide additional important information about functional hearing, particularly during natural sleep. In this study, we demonstrate how functional near-infrared spectroscopy (fNIRS) responses recorded from naturally sleeping infants might be used to estimate hearing thresholds in sleeping infants by applying a two-component response model that characterises loudness-dependent changes in haemodynamic response morphology.

fNIRS is an increasingly used neuroimaging technique for investigating auditory cortical function in infants (Lloyd-Fox et. al, 2019, Nakano et. al.,2009). fNIRS is well suited to infant studies because it is silent, portable, relatively low cost, and well tolerated, with lower susceptibility to electrical artefacts than electroencephalography (EEG)-based approaches. By measuring changes in the concentrations of oxygenated (HbO) and deoxygenated haemoglobin (HbR) associated with neurovascular coupling, fNIRS provides an indirect measure of local cortical activity (Ferrari et. al., 2012, Villringer and Chance, 1997). Numerous studies have demonstrated stimulus-evoked fNIRS responses to speech and other auditory stimuli in infants, supporting its potential for investigating auditory processing during early development (Lee et al., 2025; Lee et al., 2023; Lee et al., 2024; Mao et al., 2021; Quaresima et al., 2012; Vannasing et al., 2016).

A major challenge in extending fNIRS methodology from detection of auditory cortical responses evoked by sounds to the estimation of hearing thresholds is determining whether a response is reliably present at a given sound level. In conventional fNIRS analyses, a response is often inferred from the magnitude or statistical significance of the haemodynamic change following stimulus presentation. These approaches generally rely on assumptions about the expected temporal characteristics of the haemodynamic response and require a criterion for distinguishing stimulus-related responses from spontaneous physiological fluctuations and measurement noise (Tak and Ye, 2014, Huppert, 2016). Such assumptions may be particularly problematic in sleeping infants, in whom the morphology and magnitude of auditory-evoked fNIRS responses can vary substantially.

Our previous work (Lee et al., 2024) demonstrated that auditory-evoked fNIRS responses in naturally sleeping infants do not always conform to the canonical response morphology commonly described in infant and adult fNIRS studies (Lee et al., 2025; Lee et al., 2024). Rather than showing a consistent increase in HbO followed by a return to baseline, responses could exhibit substantial temporal variability and prolonged changes extending beyond the conventional analysis window. This variability is likely to reflect the complex physiological state of the sleeping infant and highlights the limitations of applying conventional response- inference frameworks developed primarily from adult or awake data. Importantly, these observations suggest that response detection in sleeping infants should not depend solely on whether an observed waveform conforms to a predefined canonical shape.

An alternative approach is to characterise the distribution of response measures across repeated stimulus presentations and identify statistically distinct populations corresponding to stimulus- related responses and non-responses. In our previous analysis of auditory fNIRS data from sleeping infants, we found evidence that the observed haemodynamic response could reflect the combined effects of two underlying physiological processes, an auditory canonical response and a brain arousal response (Lee et al., 2024). This motivated the development of a two-responses modelling approach in which the distribution of a response metric can be represented as a mixture of two populations. Rather than imposing a fixed response criterion based on the absolute magnitude or shape of the haemodynamic response, this approach provides a data-driven means of identifying a transition between response and non-response distributions.

In the present study, we apply a two-component-responses modelling approach to fNIRS recordings obtained from naturally sleeping infants presented with auditory stimuli at different sound intensities. We investigate whether the distribution of fNIRS response measures can be modelled and used to estimate an individual auditory hearing threshold. By modelling the relationship between the fNIRS signal and the sound level, we aim to determine whether fNIRS can provide an objective estimate of auditory sensitivity in sleeping infants. This approach may provide a foundation for extending fNIRS-based auditory assessment beyond detection of cortical activation towards modelling the changes over sound intensity to predict hearing thresholds during natural sleep.

## 2 Material and Methods

### 2.1 Participants

Twelve infants aged between 3 and 13 months participated in the study. All infants had no known history of hearing loss and had either passed newborn hearing screening or subsequent diagnostic audiological assessment. On the day of testing, all infants passed otoscopic examination and tympanometry (226-Hz or 1-kHz probe tone, as appropriate for age). Parents reported no concerns regarding their child’s general health and provided written informed consent for participation. The study was conducted in accordance with the Declaration of Helsinki and was approved by the Human Research Ethics Committee of the Royal Children’s Hospital (Project No. 71941).

### 2.2 Experiment

The experiment began with a 5-min silence baseline, during which no auditory stimuli were presented. Following the baseline period, participants were presented with blocks of repeated speech sounds at different intensities. The 12 participants were divided into two test groups (five and seven participants, respectively), and each group completed ten repetitions of its assigned set of sound levels.

Each intensity condition consisted of a 5.4-s stimulus block containing nine consecutive “ba” tokens. The speech token was a natural recording of the consonant–vowel syllable “ba” produced by a native Australian English-speaking female in a soundproof room. Each 0.45-s token was trimmed, and its mean fundamental frequency (F0) equalised using Praat (Boersma & Van Heuven, 2001). Nine token variations were then generated by shifting the F0 by ±1 Hz and varying the intensity by ±2 dB SPL. These tokens were concatenated to form each 5.4-s stimulus block. Each stimulus block was followed by 22–32 s of silence to allow the hemodynamic response to return towards baseline before presentation of the next block.

The intensity levels differed slightly between the two groups. Group A was presented with four levels (35, 50, 65, and 80 dB SPL), while Group B was presented with four levels (45, 60, 75, and 90 dB SPL). Within each set of four blocks, the order of intensity levels was pseudo- randomised. The complete stimulus presentation protocol is illustrated in Figure 1.

**Figure 1.**
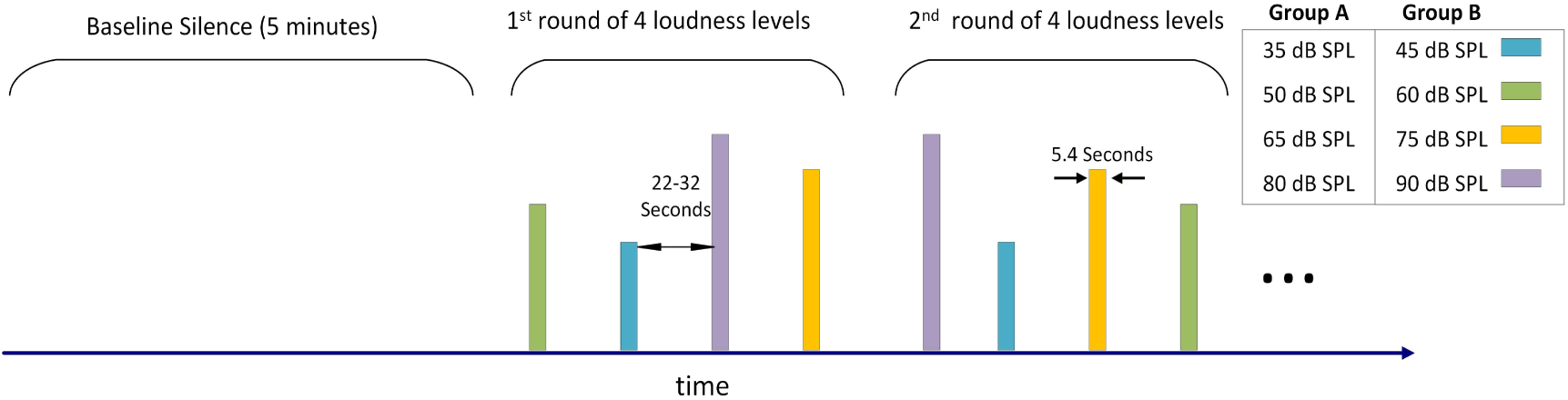
Experimental design for Cohort A and Cohort B. The experiment started with 5 minutes of silence followed by 8 stimulus blocks. Each stimulus block consisted of a 5.4- second sequence of nine “ba” tokens. Consecutive stimuli were separated by 22–32 second of silence to allow the hemodynamic response to return toward baseline. Each test completed eight repetitions of its sound intensity set (35, 50, 65, and 80 dB SPL for Group A, and 45, 60, 75 and 90 dB SPL for Group B).

### 2.3 fNIRS Data Acquisition

Infant fNIRS data were acquired using a 64-channel continuous-wave NIRScout system (NIRx Medical Technologies, LLC) equipped with flat LED optodes. The system used two wavelengths of near-infrared light (760 and 850 nm). Eight sources and eight detectors were used, yielding 18 measurement channels for each wavelength. The channels were arranged in a montage designed to cover the auditory and prefrontal regions of both hemispheres (Figure 2).

**Figure 2.**
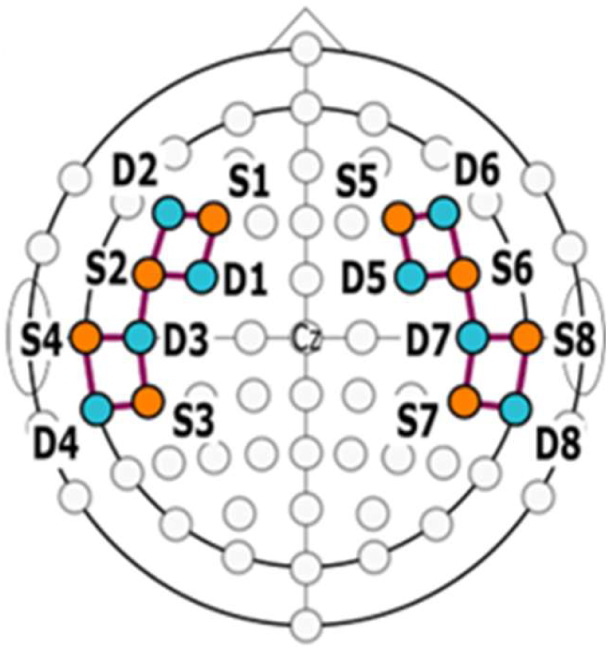
fNIRS optode montage of sources (orange) and detectors (blue) used in the study. Sources and detectors forming each ROI were as follows: Left prefrontal S1, S2, D1, D2; Left Temporal S3, S4, D3, D4; Right prefrontal S5, S6, D5, D6; Right Temporal S7, S8, D7, D8.

Short-separation channels were not included because of the practical difficulty of placing source–detector pairs at the 0.2-cm separation considered appropriate for infants. Although short-separation measurements can help characterise superficial haemodynamic signals, previous infant fNIRS work found that removing short-channel contributions did not alter the statistical inferences drawn from the data, suggesting that the benefit of short-channel correction may be task- and context-dependent in this population (Brigadoi & Cooper, 2015; Emberson et al., 2016).

### 2.4 Data Processing

All fNIRS data preprocessing was performed using the MNE-Python toolbox. The raw light intensity data were first converted to optical density. Signal quality was evaluated for each channel using the Scalp Coupling Index (SCI) function (Pollonini et al., 2014), which calculates the correlation between signals measured at the same source–detector pair but at different wavelengths. A low correlation coefficient indicates poor optode–scalp contact; in this study, channels with an SCI below 0.8 were considered to have poor signal quality and were discarded. Overall, 5.6% of channels had an SCI below this threshold and were excluded from further analysis.

Motion artefacts and physiological noise were corrected using Temporal Derivative Distribution Repair (TDDR; Fishburn et al., 2019), followed by band-pass filtering using a Butterworth filter with cutoff frequencies of 0.01 and 0.25 Hz. The processed optical density data were then converted to oxy- and deoxy-haemoglobin (HbO and HbR, respectively) concentration changes using the modified Beer–Lambert law with a partial pathlength factor of 0.1.

The data were epoched from -3.0 s to 27.4 s relative to stimulus onset. Baseline correction was performed by subtracting the mean haemoglobin concentration change during the -3.0 s to 0 s pre-stimulus period from each epoch. Epochs containing extreme values exceeding ±80 mmho were considered as containing noise and were rejected from further analysis.

### 2.5 Speech threshold for infants with normal hearing

Traditional speech thresholds often use highly audible, two-syllable “spondee” words (e.g., baseball, hotdog). Consonant-Vowel (CV) words have significantly lower acoustic energy and brief durations. As a result, the physical threshold to detect a brief CV syllable can sit slightly higher than long words—closer to 13 to 18 dB SPL, depending on whether the starting consonant is a high-energy sonorant (e.g., “ma”) or a low-energy voiceless fricative (e.g., “pa” or “ta”). The standard deviation for speech threshold testing in healthy populations is roughly 3 to 5 dB. In our study, CV syllable “Ba” was used; therefore, the average normal hearing threshold is approximately 18 dB SPL (equivalent approximately to 0 dB HL) with a SD of 5 dB SPL, i.e., a confidence interval of 8 to 28 dB SPL, which corresponds to roughly -10 to 10 dB HL on a clinically calibrated scale.

### 2.6 Two-responses fNIRS Model

In our previous studies, we demonstrated that speech-evoked fNIRS responses in naturally sleeping infants consist of two concurrent response components (Lee et al., 2024; Lee et al., 2025). Lee et al. (2024) investigated the morphology of the HbO response to detection of the same 5.4-s “ba” stimulus at 65 dB SPL. Using independent component analysis, they showed that the response was composed of a positive going HbO response peaking at 6-7 s after stimulus onset and a negative-going HbO response peaking at 15-17 s after stimulus onset. They proposed that the positive-going component, with a standard canonical shape, was the auditory response and the negative going response was associated with activation of arousal mechanisms. Over repeated trials, the negative component habituated rapidly (as expected for an arousal response) altering the overall morphology as the test progressed. Lee et al. (2025) applied these 2 component findings to the morphology of the speech sound discrimination response. Both weak (“ba” vs “ga”) and strong (“ba” vs “bee” and “tea”) contrasts were tested. Two main findings were that the negative arousal component was much weaker for the very subtle contrast (“ba“vs “ga”) compared to the strong contrasts, and that the positive auditory response component increased in latency for the subtle contrast. We therefore hypothesised that the effects of stimulus intensity for the detection response would follow the same patterns: i.e. the negative components would decrease in amplitude, and the positive response would increase in latency, as the intensity was lowered. Furthermore, we hypothesised that the negative relative to positive amplitude ratio would decrease as the number of trials increased, due to the negative response habituating more rapidly than the positive response. Based on these hypotheses, we developed a model to describe how the HbO response morphology would change across different sound intensities for normal hearing infants.

To simplify our model for the purpose of this proof-of-principle demonstration, we modelled the two components as gaussian shapes and had only two parameters that changed with intensity: the latency of the positive component and the relative amplitude ratio of the negative versus positive peak amplitudes. We expect both component absolute amplitudes to be affected by the same infant-specific factors (such as position of optodes and thickness of skull for example) but hypothesised that the relative amplitude would decrease as the number of trials increased and/or the intensity decreased. Thus, different models of relative amplitudes are needed depending on the number of trials analysed in the experiment. In the results section we developed and used the model for the first 4 trials of each sound intensity and for temporal and prefrontal regions separately.

For this proof-of-principle exercise, we first estimated a small number of model parameter values based on data from Lee et al (2024) for the 65 dB SPL intensity Specifically, we fixed the latency of the negative component at 15 s, and we estimated the latency range of the positive component to between 5.4 s to 20 s. We then constructed heuristic functions that related each parameter to intensity while passing through the determined parameter values. For the parameter of negative/positive amplitudes we set this based on the Lee et al. (2024) paper at 0.8 and set the parameter range to (0, 6). This procedure enabled a template morphology to be determined for any arbitrary intensity. The templates represent the morphology to be expected for an infant with normal hearing listening to that intensity. Alternatively, for an infant with unknown hearing levels, the best template fit represents the equivalent loudness at which the infant heard the stimulus, using the assumption that it is the perceived loudness rather than the physical sound intensity that drives the morphology changes at a cortical level (Weder et al., 2020). For example, if a 65 dB SPL sound elicited a response that best fit the morphology for 40 dB SPL in normal hearing infants, it means that the infant heard the sound with the same loudness that normal hearing infants hear the 40 dB SPL stimulus. In this paper we denote this loudness as “equivalent normal loudness” or ENL.

### 2.7 Hearing thresholds seeking procedures

Using the derived template responses, we implemented a general threshold prediction framework. With steps as follows:

1. For each stimulus intensity, the recorded response is compared with a range of normal- hearing templates corresponding to different stimulus intensities to quantify the likelihood of correspondence to each template. The resulting distribution of likelihood values is then summarised using the interquartile range (IQR), which characterises the spread of the likelihood distribution and provides an indication of the consistency of the template-matching results. These metrics characterise the degree and consistency of alignment between the observed and template responses across sound intensity levels.
2. For each intensity in an experiment (and each region of interest), the best fit template intensity or ENL and interquartile range are plotted against the actual stimulus intensity. A curve-fitting procedure based on Monte Carlo sampling is then used to generate a distribution of plausible linear regression lines. Each fit was weighted according to the interquartile range (IQR) at each stimulus level, with larger IQRs indicating greater response variability and therefore lower measurement reliability. Accordingly, stimulus levels with larger IQRs were assigned lower weights in the regression, giving greater influence on data from stimulus levels with more consistent HbO responses. Stimulus levels generating an IQR greater than 40 dB SPL were considered to indicate poor fNIRS signal quality and were excluded from the hearing-threshold prediction model. i.e., the weighting is set to zero if IQR is greater than 40 dB SPL. A linear fit of ENL versus intensity is expected from psychophysical experiments plotting loudness versus intensity, with participants with sensory loss also exhibiting a linear relationship but greater slope due to recruitment.
3. The threshold and its standard deviation are determined from the intersection of the regression lines with the threshold loudness (ENL of 18 dB SPL).

An example of the hearing threshold estimation steps is shown in Figure 3. The top row illustrates the fit between the recorded responses at 80 dB SPL and the normal-hearing templates (Step 1). Solid lines represent the recorded HbO responses (average of 4 epochs) in the prefrontal and temporal regions, while dotted lines indicate the best-fitting normal-hearing templates. Under the averaged waveforms, the bars represent how the MSE (mean squared error) of template fits to the data vary with template intensity (or ENL - equivalent normal loudness). In this example, the recorded responses in the prefrontal and temporal regions yield the highest likelihoods for ENLs of 80 dB SPL and 90 dB SPL, respectively (shown in red). The bottom row (illustrating Step 2) shows the best fit template intensity and interquartile range for each sound intensity. The thick bars indicate the interquartile range (IQR; Q1–Q3) of the likelihood distribution, with thin tails marking the lower (Q1) and upper (Q3) quartiles. The shaded area shows the distribution of plausible linear regression lines. The horizontal red bar indicates the speech threshold for normal hearing infants, purple bar illustrates Step 3, showing the predicted threshold in dB SPL.

**Figure 3.**
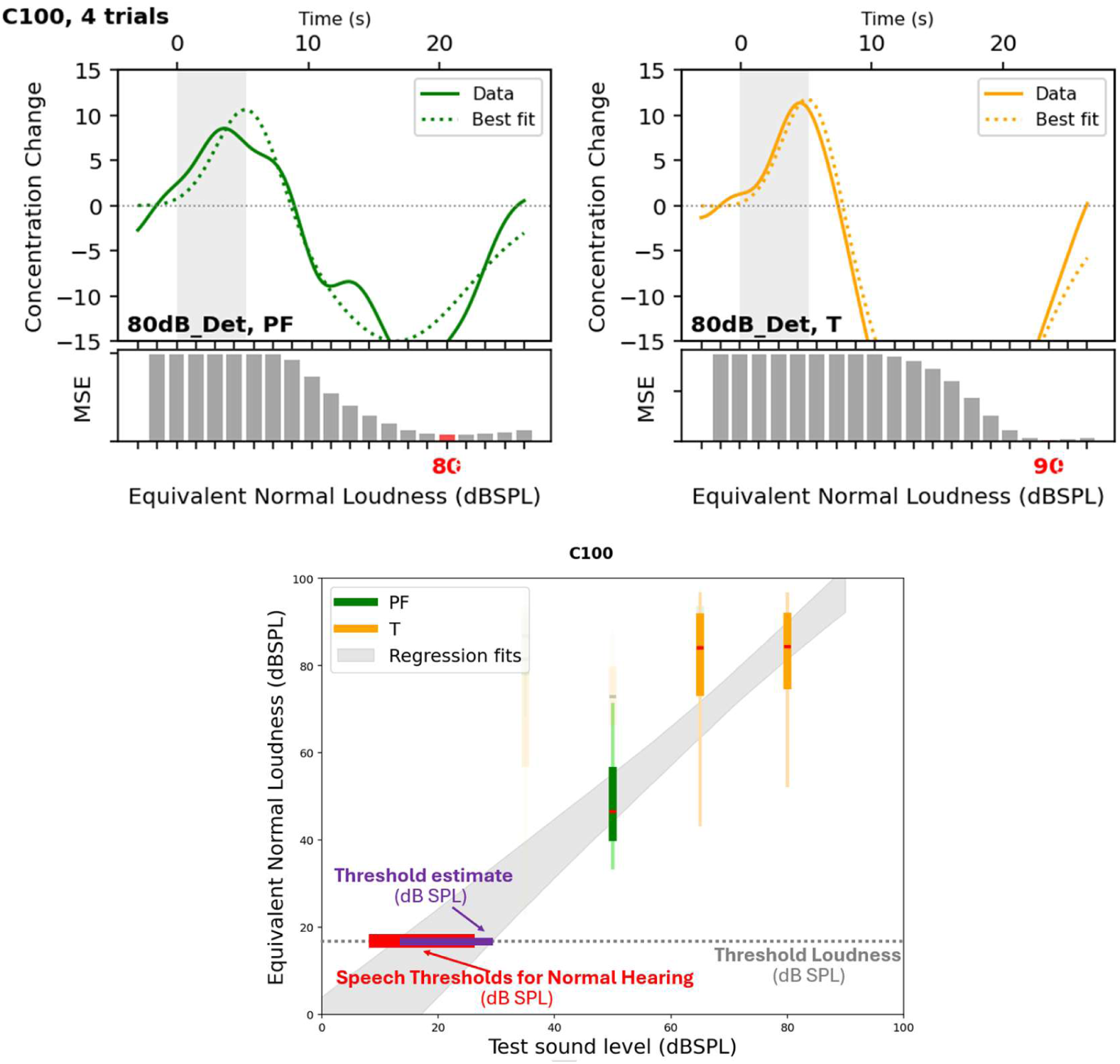
An example of the hearing threshold estimation procedure is shown in Fig. 3. The top row illustrates the fit between the recorded responses at 80 dB SPL and the normal-hearing templates. The bottom row shows the best fit template intensity and interquartile range for each sound intensity and the distribution of plausible linear regression lines.

## 3 Results

### 3.1 Responses of normal hearing infants

Figure 4 shows the grand average HbO and HbR responses for the two test groups of infants averaged across regions of interest. The grand average was calculated for 4 trials. Higher amplitudes were observed for both the positive and negative components at test sound intensities above 65 dB SPL in both groups.

**Figure 4.**
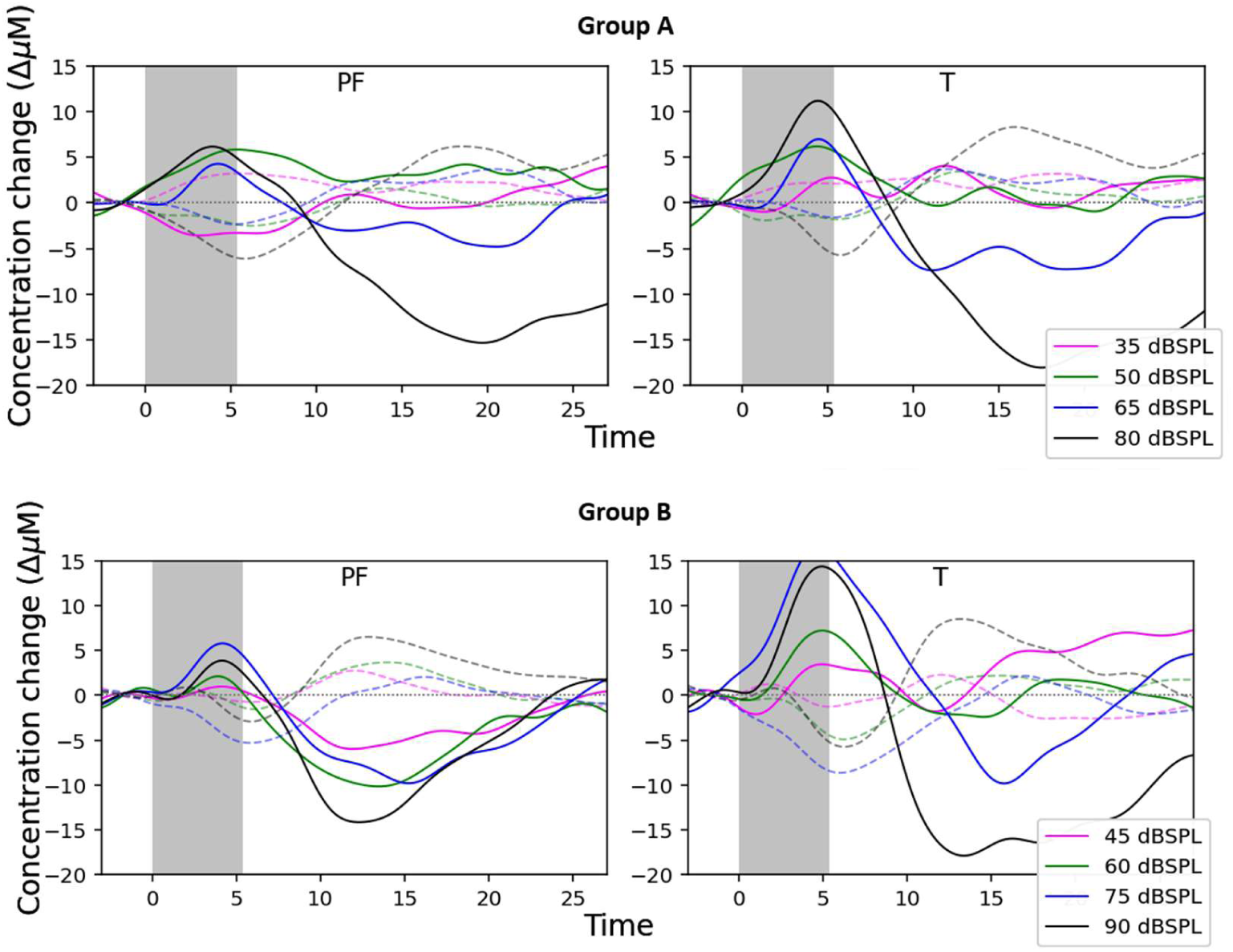
Grant average HbO (solid lines) and HbR (dotted lines) responses for the two groups averaged across pre-front and temporal regions.

### 3.2 Responses of normal hearing infants

The infants in this experiment had presumed normal hearing. However, they are likely to have a range of actual hearing thresholds. Additionally, although they have passed newborn screening, and normal tympanometry, these tests do not exclude the possibility of a mild hearing loss. As a first step here however, we use our threshold seeking procedure to estimate hearing threshold for each of the 12 infants individually. A total of 2,000 resampling iterations were performed to generate a distribution of plausible linear fits for each infant. when estimating the hearing threshold.

Figure 5 shows the constrained Monte Carlo regression fits for Groups A and B. For each infant, the green bars represent the IQR of the prefrontal HbO response at each stimulus level, while the thick orange bars represent the IQR of the temporal HbO response. The regression model was used to characterise the relationship between HbO response and sound intensity, with the resulting distribution of plausible linear regression lines represented by the shaded area. Stimulus levels with an IQR greater than 40 dB SPL were considered to indicate poor fNIRS signal quality and were excluded from the hearing-threshold prediction model; the corresponding IQR bars are therefore shown with reduced brightness to indicate that these data were excluded from the Monte Carlo regression.

**Figure 5.**
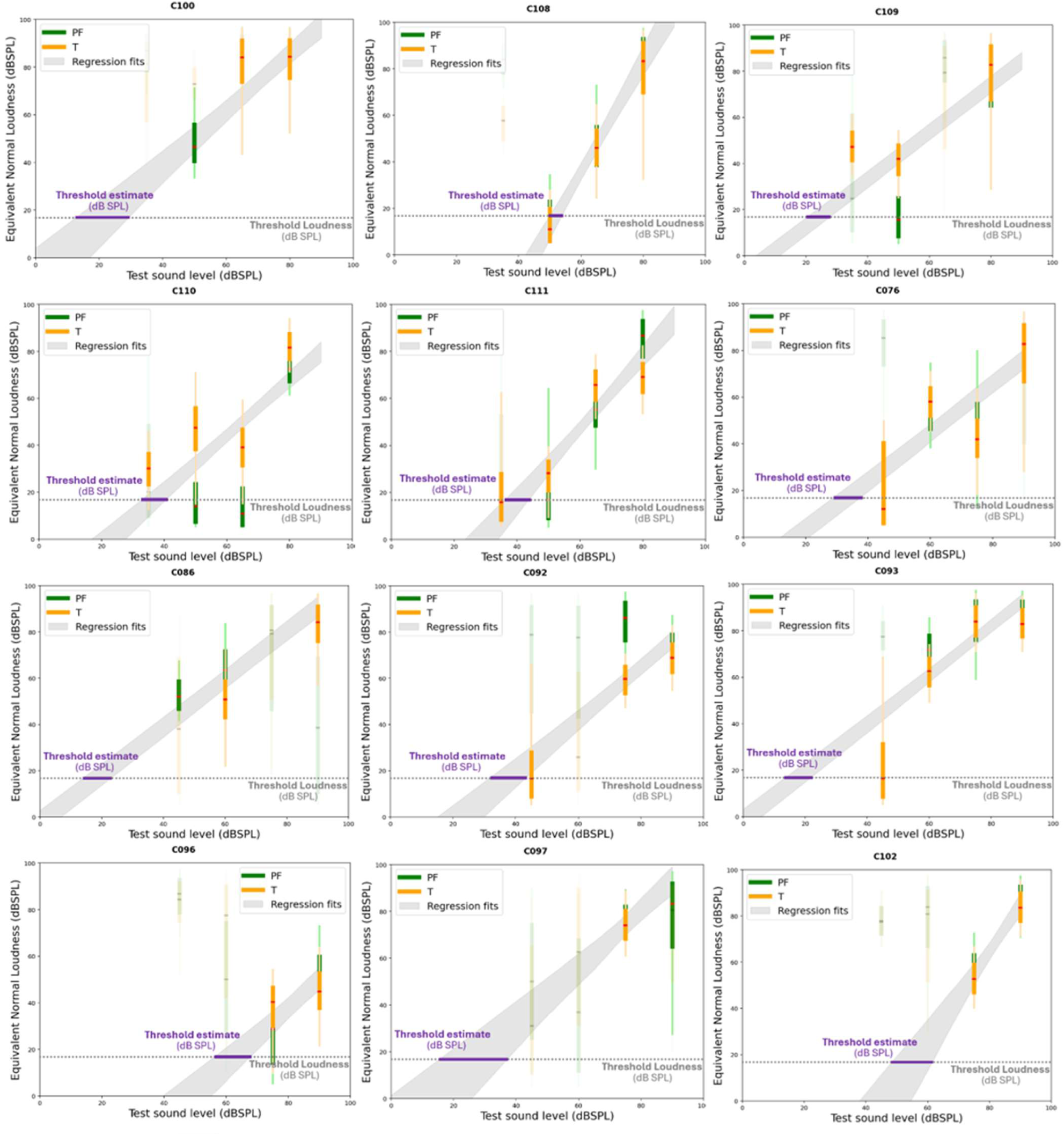
Equivalent Normal Loudness estimation for both test groups. The orange and green bars show the best fit template intensity and interquartile range for each sound intensity in the pre-front and temporal regions, respectively. The shaded area shows the distribution of plausible linear regression lines, the horizonal purple bar indicates the estimated threshold.

Across the 12 infants, 9 had predicted hearing thresholds within or close to the expected range for normal hearing (-10 to 10 dB HL), while three infants (C108, C096, and C102) had predicted thresholds above 35 dB HL. Notably, all three infants with higher predicted thresholds lacked valid fNIRS responses from both temporal and prefrontal regions at the lowest sound intensity tested, i.e., those responses were excluded from the prediction model because they exceeded the predefined IQR criterion for acceptable signal quality. Consequently, the regression models for these infants were based on fewer reliable observations, particularly at the lower sound intensities, which likely contributed to the elevated predicted thresholds.

Overall, the mean predicted hearing threshold across the 12 infants was 17.13 dB HL with a standard deviation of 14.42 dB HL. When the three infants with invalid fNIRS responses at the lowest sound intensity were excluded, excluding the 3 infants with invalid fNIRS recording at the lowest sound intensity level, the mean predicted threshold across the 9 infants was 10.72 dB HL, with a standard deviation of 8.64 dB HL.

### 3.3. Case study: ENL estimation in hearing impaired infants

We estimated the ENL for a subject with hearing impairment. Subject C101 was a 7-month- old infant with bilateral mild to moderate hearing loss, with ABR thresholds of 40 dBnHL for both ears. The fNIRS study was conducted via tubephone to the right ear. The fNIRS signal was processed and the ENL was estimated by following the same protocols as the normal hearing groups. Figure 6 shows the estimated ENL for subject C101.

**Figure 6.**
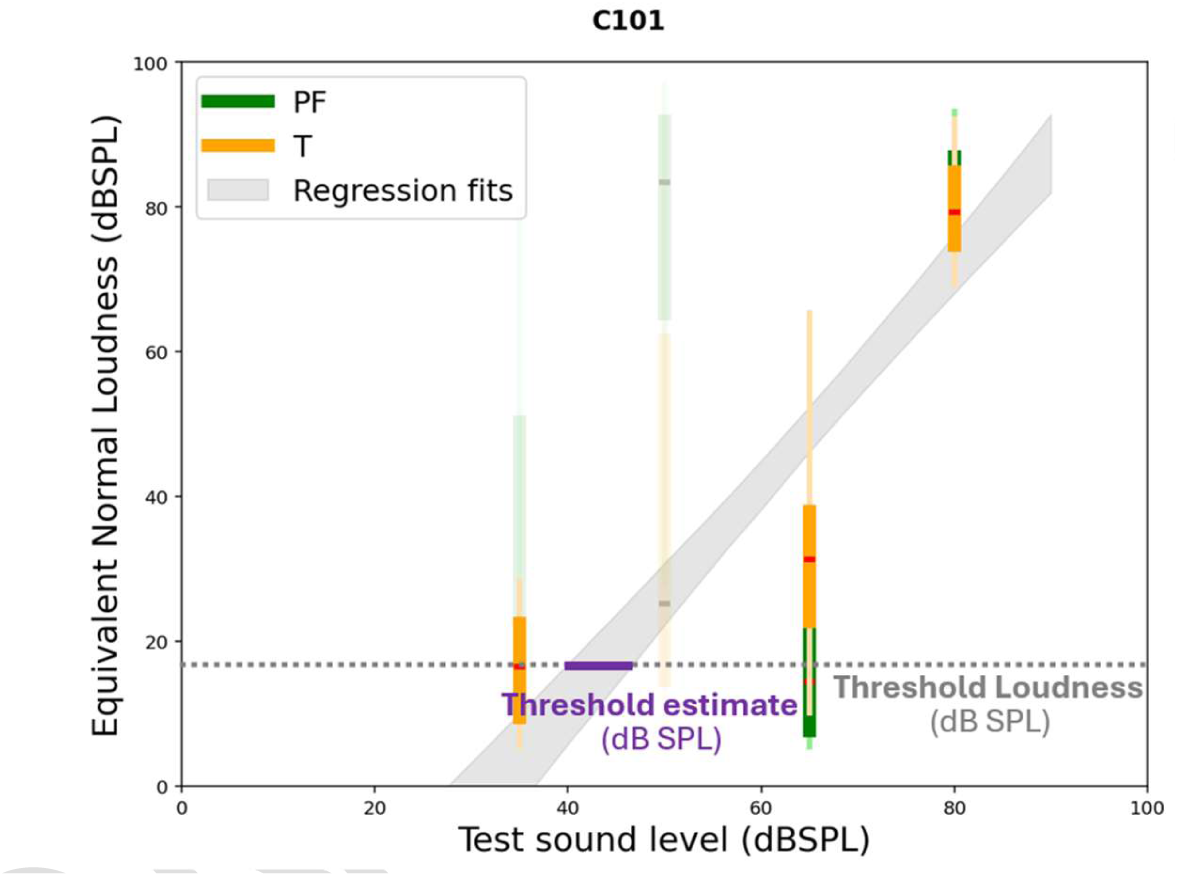
Equivalent Normal Loudness estimation for subject C101. The orange and green bars show the best fit template intensity and interquartile range for each sound intensity in the pre-front and temporal regions, respectively. The shaded area shows the distribution of plausible linear regression lines, the horizonal purple bar indicates the estimated threshold.

Figure 6 shows the ENL estimates for subject C101. The estimated ENL ranges from 21 to 27 dB HL, indicating mild hearing loss. We note that three of the four IQR at test levels of 35 and 50 dB SPL exceed 40 dB SPL. This variability may reflect low fNIRS signal quality or unstable fNIRS responses, potentially because these test levels are close to the participant’s hearing thresholds.

We also note that, in this case study, the regression lines were estimated using the same curve- fitting procedure as in the normal-hearing group, namely linear regression. This approach was used because the participant was tested without hearing aids. However, for aided infants, alternative regression models may be more appropriate to account for potentially non-linear loudness growth functions.

## 4 Discussion

### 4.1 Hearing threshold prediction in normal hearing infants

A key finding of this study was that the proposed fNIRS-based modelling approach produced hearing-threshold estimates within the expected range for normal hearing (-10 to 10 dB HL) in 9 of the 12 infants (75%). Over the 9 infants, the mean predicted threshold was 10.72 ± 8.64 dB HL, indicating that the model generally produced threshold estimates within or close to the expected normal-hearing range. These findings provide preliminary evidence that haemodynamic responses measured using fNIRS can be quantitatively related to sound intensity and used to derive an objective estimate of auditory sensitivity in infants. Importantly, the predicted thresholds were derived from the relationship between the HbO response and stimulus intensity rather than from a direct behavioural response, suggesting a potential approach for estimating hearing sensitivity in infants who may be unable to provide reliable behavioural responses.

Some variability in predicted thresholds was evident across infants, which is expected given the substantial inter-individual variability associated with infant fNIRS measurements. Factors may include differences in cortical haemodynamic responses, head and optode positioning, individual scalp and skull anatomy. The constrained Monte Carlo regression framework used in this study provides a means of accounting for this variability by incorporating the uncertainty in the fitted response–intensity relationship and the resulting threshold estimates. Despite these sources of variability, the threshold estimates for most of the infants were within the expected normal-hearing range, suggesting that the relationship between sound intensity and cortical haemodynamic response can be used for threshold estimation.

Over all infants, the mean predicted threshold was 17.13 ± 14.42 dB HL. Three infants with predicted thresholds above 35 dB HL require particular consideration. In these infants, the fNIRS response at the lowest stimulus intensity was considered unreliable according to the predefined IQR criteria and was therefore excluded from the regression model. Consequently, the model had fewer observations with which to characterise the response–intensity relationship, particularly at the lower end of the stimulus range. This may have contributed to greater uncertainty in the extrapolated threshold estimates. These findings highlight the importance of obtaining reliable fNIRS responses at low stimulus intensities for accurate estimation of hearing thresholds.

The variability observed in the predicted thresholds was comparable to that reported for standard clinical hearing-threshold measurements (Primus, 1991, Widen et. al., 2000, Parry et. al., 2003). This comparison is relevant because some degree of variability is inherent in hearing-threshold estimation, particularly in infants, and an exact correspondence between fNIRS-derived and clinical thresholds may therefore not be expected. Rather, the present results suggest that fNIRS can provide threshold estimates with clinically plausible variability while using a non-invasive objective measure.

An additional strength of the present modelling framework is its incorporation of response variability into the threshold estimation process. By weighting observations according to the IQR of the HbO response, measurements with greater variability were given less influence on the estimated response–intensity relationship. This provides a principled approach to handling the substantial variability inherent in infant fNIRS data. Our results also suggest that improving fNIRS signal quality and increasing the reliability of cortical responses at lower sound intensities will improve the accuracy and robustness of the approach.

### 4.2 Method usability in hearing evaluation in hearing impaired infants

#### 4.2.1 Unaided hearing thresholds in hearing impaired infants

The case study of subject C101 provides preliminary insight into the potential applicability of the proposed fNIRS-based modelling approach to hearing evaluation in infants with hearing impairment. The estimated ENL ranged from 21 to 27 dB HL, suggesting mild hearing loss. Although this single case does not establish the accuracy of the method in hearing-impaired infants, it demonstrates the feasibility of applying the proposed framework to estimate auditory sensitivity beyond the normal-hearing group. This potential is particularly important for estimating hearing thresholds in infants with auditory neuropathy, where existing methods based on electrophysiology are not feasible.

Several methodological considerations are important when extending this approach to hearing- impaired infants. In the case study, substantial variability was observed in the fNIRS responses at some stimulus intensities, as reflected by the large interquartile ranges. This variability may arise from low fNIRS signal quality or less stable cortical responses, particularly when stimulus levels are close to the participant’s hearing threshold. As observed in the normal-hearing group, unreliable responses at lower stimulus intensities may affect the fitted response–intensity relationship and increase uncertainty in the resulting threshold estimates. Improving signal quality and obtaining reliable responses across a wider range of stimulus intensities will therefore be important for improving the robustness of threshold estimation.

Another consideration is the choice of regression model. In the present case study, linear regression was used, following the same curve-fitting procedure as in the normal-hearing group. However, hearing-impaired infants may have a reduced listening dynamic range, potentially resulting in a different relationship between sound intensity and the cortical haemodynamic response (Moore, 1995, Marozeau and Florentine, 2007). Although loudness growth functions can be non-linear close to hearing threshold (Marozeau and Florentine, 2007), in this study we assumed that a linear model was appropraite for both normal-hearing and hearing-impaired infants. This assumption will not hold for aided infants, where the loudness growth will be influenced by the input/output function of the hearing aid, which may not be linear. Alternative regression models that better accommodate changes in response patterns across stimulus intensities may be more appropriate for this population. Future studies involving larger cohorts of hearing-impaired infants are needed to investigate the suitability of different regression models and establish their performance against established hearing assessment methods.

#### 4.2.2 Verification of hearing aid and cochlear implant program parameters

Regardless of the shape of the aided loudness growth functions, our estimates of the aided ENL enable estimates of whether the aid has been adequately programmed so that important speech elements are clearly audible to the infant and fit within their aided perceptual dynamic range. For example, any hearing device should aim to amplify conversational level speech to a perception loudness similar to that which normal hearing infants would perceive. The potential use of ENL estimates is particularly important for cochlear implant programming verification, where setting MAP parameters and verification usually depends solely on unreliable behavioural observations.

## 5 Conclusion

This work demonstrates the feasibility of using fNIRS and HbO-based modelling to estimate auditory sensitivity in infants. In the normal-hearing group, 75% of infants produced threshold estimates within the expected normal-hearing range (an estimated mean ENL of 10.27 dB HL, with a standard deviation of 8.64 dB HL), providing encouraging preliminary evidence that cortical haemodynamic responses can be quantitatively related to sound intensity for hearing threshold estimation. The case study further illustrates the potential applicability of this approach to infants with hearing impairment, while highlighting the importance of reliable fNIRS responses at lower stimulus intensities and appropriate regression model selection. Although these findings are promising, the small sample size and variability in threshold estimates warrant cautious interpretation. Further studies involving larger cohorts and infants with a broader range of clinically established hearing thresholds are needed to evaluate the accuracy, reliability, and clinical utility of the proposed approach. With further validation and methodological refinement, fNIRS-based modelling may provide a valuable objective complement to existing hearing assessment methods, particularly for infants who cannot provide reliable behavioural responses.

## 6 Acknowledgements

The authors disclosed receipt of the following financial support for the research, authorship of this article: CASS medicine and science grant, Garnett Passe and Rodney Williams Memorial Foundation, Victorian Government Medical Research Acceleration Fund, National Health and Medical Research Council. Demi Gao is supported by Garnett Passe and Rodney Williams Memorial Foundation Mid-Career Fellowship. The Bionics Institute acknowledges the support it receives from the Victorian Government through its Operational Infrastructure Support Program.

## 8 Data availability

The data that support the findings of this study are available from the C. M. McKay, upon reasonable request.

